# Lung Cancer Explorer (LCE): an open web portal to explore gene expression and clinical associations in lung cancer

**DOI:** 10.1101/271056

**Authors:** Ling Cai, ShinYi Lin, Yunyun Zhou, Lin Yang, Bo Ci, Qinbo Zhou, Danni Luo, Bo Yao, Hao Tang, Jeffrey Allen, Kenneth Huffman, Adi Gazdar, John Heymach, Ignacio Wistuba, Guanghua Xiao, John Minna, Yang Xie

## Abstract

We constructed a lung cancer-specific database housing expression data and clinical data from over 6,700 patients in 56 studies. Expression data from 23 “whole-genome” based platforms were carefully processed and quality controlled, whereas clinical data were standardized and rigorously curated. Empowered by this lung cancer database, we created an open access web resource – the Lung Cancer Explorer (LCE), which enables researchers and clinicians to explore these data and perform analyses. Users can perform meta-analyses on LCE to gain a quick overview of the results on tumor vs normal differential gene expression and expression-survival association. Individual dataset-based survival analysis, comparative analysis, and correlation analysis are also provided with flexible options to allow for customized analyses from the user.

## Introduction

Lung cancer is the leading cause of cancer-related death worldwide. Despite tremendous efforts put toward diagnosis and treatment, the five-year survival rate of lung cancer is still as low as 18%[1]. Over the past few decades, advancements in genome profiling techniques have greatly improved our understanding of cancer development at the molecular level, and have enabled the discovery of biomarkers that facilitate individualized cancer treatments. With the advent of public data repositories of genome profiling data, such as the Gene Expression Omnibus (GEO, [2]), ArrayExpress[3, 4], and The Cancer Genomics Atlas (TCGA), it has become increasingly important and beneficial for researchers to mine the available data sets to discover potential biomarkers and test new biological hypotheses.

Despite the wealth of information offered by such data, utilization of public datasets is not easy, and often it can be prohibitively challenging. There is a plethora of lung cancer patient data published each year, but the data are scattered around in different public data depositories or at individual websites. There are often inconsistencies for the same patient cohort among different websites, likely due to differences in preprocessing approaches and the versions of platform annotations. Moreover, clinical records from different studies are often summarized using different terminologies. Proper usage of publicly available datasets requires specialized expertise in acquiring, processing, normalizing and filtering of the data, which are challenging for general researchers and clinicians. To facilitate researchers to utilize public datasets for biomarker discovery, a number of re-annotated database have been developed, including OncoMine[5], GeneSapien[6], Gemma[7], M2DB[8], CancerMA[9], cBioPortal[10], KMPlot[11], PrognoScan[12], PROGgene[13] and so forth.

In this study, we describe our development of a new web application, Lung Cancer Explorer (LCE) (http://lce.biohpc.swmed.edu/), populated by a centralized lung cancer database. Compared to other existing databases, our database houses the largest collection of lung tumor expression data from 56 studies for over 6,700 patients (Table1, Figure 1 and Supplementary Table 1). Tremendous efforts were made in manual curation and standardization of the datasets so that they could be used for meta-analysis. The user-friendly open web portal provides several easy but versatile analysis tools. These tools include meta-analysis, which enables users to gain a quick overview of the results from all datasets while combining statistical power from multiple datasets, as well as individual dataset-based analyses that allow for more flexibility and customization from the user.

**Figure 1.**
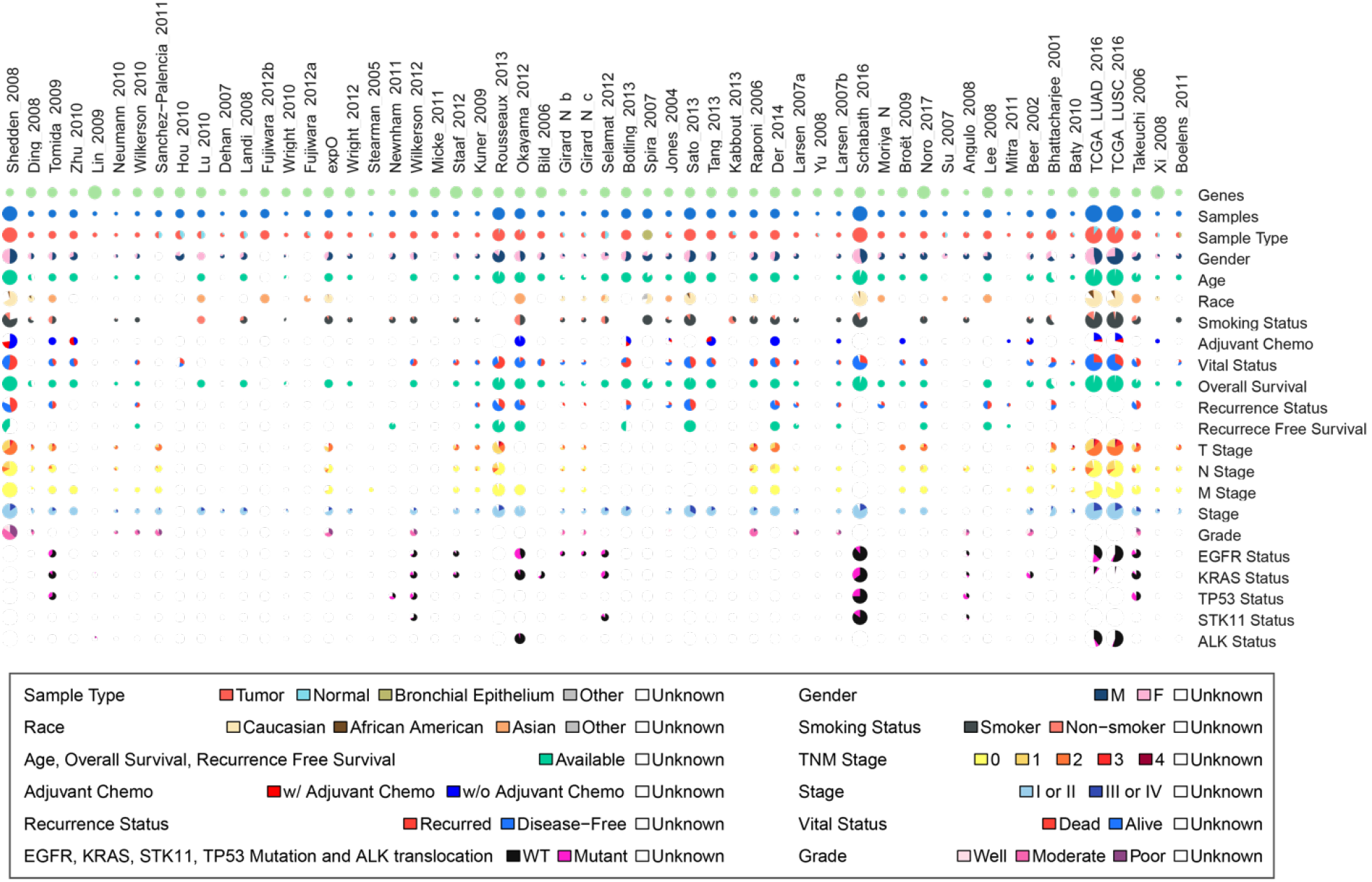
Summary of lung cancer database variable distribution. This is an overview summary of the lung cancer database that feeds into the Lung Cancer Explorer. Gene expression data and clinical data were collected from 56 studies that include over 6,700 patients. For each study and each variable, a pie chart is used to summarize the data. The size of the pie chart is proportional to the number of samples from the study, and the color scheme for the pie chart sectors are provided below the gridded pie charts. All the detailed numbers can be found in Supplementary Table 1.

## Materials and Methods

### Data collection

Datasets were collected from GEO, TCGA and individual literatures. The search from GEO was performed by GEOmetadb [14]. For datasets that have not been deposited into GEO, we made our selection through a literature search and by referencing other commonly used databases.

### Clinical data curation

Clinical data for datasets deposited into GEO were retrieved from GEO by R package GEOquery; TCGA clinical data were downloaded from Sage Bionetworks’ Synapse database [15], and other datasets were downloaded from sources provided in the original publication. The clinical data obtained directly from these public domains often contains non-standard terminology. To standardize the clinical variables from different studies, codebooks were devised for each variable in order to ensure the accuracy and compatibility of the clinical annotation from different sources (Supplementary Table 3). The patient histology codebook was created based on the 2015 World Health Organization (WHO) Classification of Lung Tumors [16] (Figure 2 and Supplementary Table 2). For the TCGA lung cancer data in particular, instead of using the histology classification provided by the patient information file, histology was determined based on expression signature as developed by Girard et al [17], as the study has shown an improved classification accuracy with the gene expression classifier on the TCGA data. Consequently, the histology-misclassified samples were excluded from the TCGA cohorts in cancer specific meta-analysis. For all datasets, programmatic and manual data curations were carried out and the procedures were repeated three times with scrutiny. For our records, all the data handling steps were saved with detailed documentation.

**Figure 2.**
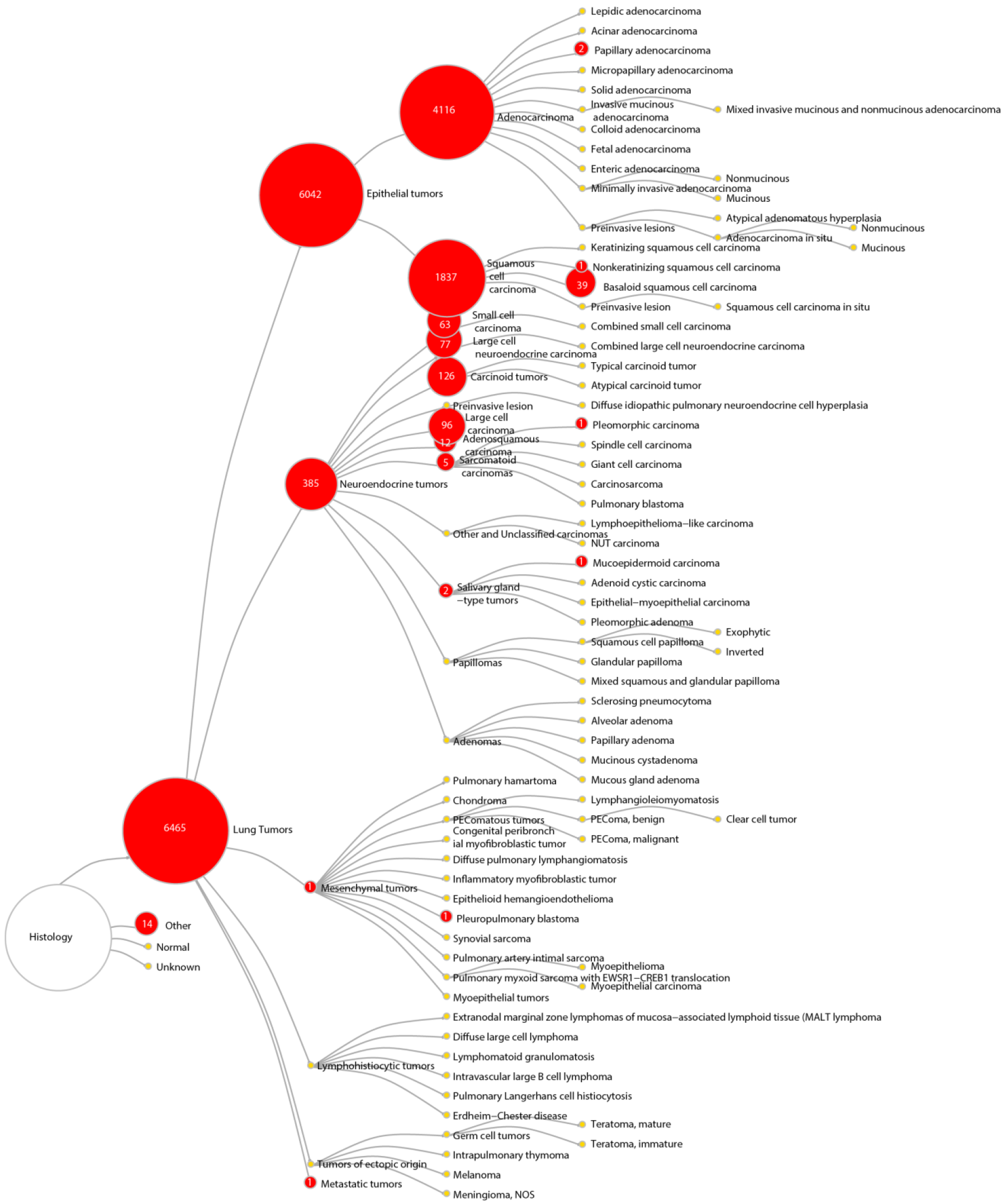
Histology classification of samples collected in the lung cancer database. This tree diagram represents the hierarchical structure of the 2015 WHO classification system of lung tumors. Numbers on the red nodes denote the number of samples from the lung cancer database belonging to the corresponding histology type.

### Expression data processing

Expression data for datasets deposited into GEO were retrieved from GEO by R package GEOquery. TCGA expression data were downloaded from Broad GDAD firehose [18], and other datasets were downloaded from sources provided in the original research paper. It is not uncommon in the field of biomarker discovery that the signatures have poor reproducibility in other data sets. Such discrepancy could be at least partially attributed to the differences in experimental settings, sample handling,
measurement platforms and, importantly, data processing procedures. The datasets collected in this study were generated from 23 different platforms, with the majority being microarrays. We adopted different strategies to process the data (Supplementary figure 1) to convert the expression data from probe level to gene level.

### Database structure/ web interface

Our web application Lung Cancer Explorer (LCE) can be accessed through http://lce.biohpc.swmed.edu/. It was created using PHP (7.0.12-1) in the R Programing environment (3.3.1) with MySQL database (Ver 14.14 Distrib 5.5.49) in the backend. Our MySQL database contains tables for samples, patients and gene expression data with supporting data dictionaries (Figure 5, Supplementary Table 3).

### Survival analysis

Survival curves were estimated using the product-limit method of Kaplan-Meier [19] (survival, R package [20]). R package *mclust [21]* was used to identify the gene expression cut-off based on Gaussian mixture model clustering, assuming a bi-modal distribution when users select the “cluster” option under the survival analysis module of LCE. A log-rank test was used to compare the survival differences among different patient groups. A Cox Proportional Hazard regression model was used to assess the survival association and calculate the hazard ratio (HR) with the continuous gene expression in each individual dataset.

### Meta-analysis

For survival meta-analysis, the R package meta [22] was used to calculate the summary HR from the HRs of individual datasets. For tumor vs normal differential expression meta-analysis, R package metafor [23] was used to calculate the summary standardized mean difference (tumor – normal) using Hedges’ G as an effect size metric.

## Results

### Construction of the Lung Cancer Database

Over a span of 5 years, we have collected 56 datasets generated by 23 “whole-genome”-based expression platforms (See “Data collection”, “Clinical data curation” and “Expression data processing” sections in Methods). The overarching goal is to include datasets with large numbers of samples as well as datasets with more comprehensive coverage of clinical information with an emphasis on survival data.

**Table 1.** Provides a summary of the sources and sample numbers of the datasets in our database.

| Friendly Name | Pubmed ID | GEO ID | Sample Number |
| --- | --- | --- | --- |
| Shedden_2008 | 18641660 |  | 442 |
| Ding_2008 | 18948947 | GSE12667 | 75 |
| Tomida_2009 | 19414676 | GSE13213 | 117 |
| Zhu_2010 | 20823422 | GSE14814 | 133 |
| Lin_2009 | 19737969;20668451 | GSE16534 | 43 |
| Neumann_2010 | 20196851 | GSE17475 | 28 |
| Wilkerson_2010 | 20643781 | GSE17710 | 51 |
| Sanchez-Palencia_2011 | 20878980 | GSE18842 | 91 |
| Hou_2010 | 20421987 | GSE19188 | 156 |
| Lu_2010 | 20802022 | GSE19804 | 120 |
| Dehan_2007 | 17258348 | GSE1987 | 37 |
| Landi_2008 | 18297132 | GSE10072 | 107 |
| Fujiwara_2012b | 21737174 | GSE20853 | 164 |
| Wright_2010 | 20544843 | GSE20875 | 36 |
| Fujiwara_2012a | 21737174 | GSE2088 | 87 |
| expO |  | GSE2109 | 141 |
| Wright_2012 | 22514692 | GSE23822 | 56 |
| Stearman_2005 | 16314486 | GSE2514 | 39 |
| Newnham_2011 | 21385341 | GSE25326 | 85 |
| Wilkerson_2012 | 22590557 | GSE26939 | 102 |
| Micke_2011 | 22011649 | GSE28571 | 100 |
| Staaf_2012 | 22676229 | GSE29016 | 72 |
| Kuner_2009 | 18486272 | GSE10245 | 48 |
| Rousseaux_2013 | 23698379 | GSE30219 | 307 |
| Okayama_2012 | 22080568;23028479 | GSE31210 | 224 |
| Bild_2006 | 16273092 | GSE3141 | 111 |
| Girard_N_b |  | GSE31547 | 50 |
| Girard_N_c |  | GSE31548 | 50 |
| Selamat_2012 | 22613842 | GSE32863 | 116 |
| Botling_2013 | 23032747 | GSE37745 | 196 |
| Spira_2007 | 17334370;20375364 | GSE4115 | 192 |
| Jones_2004 | 15016488;21737174 | GSE1037 | 80 |
| Sato_2013 | 23449933;24850841;27354471 | GSE41271 | 275 |
| Tang_2013 | 23357979 | GSE42127 | 176 |
| Kabbout_2013 | 23659968 | GSE43458 | 110 |
| Raponi_2006 | 16885343 | GSE4573 | 130 |
| Der_2014 | 24305008 | GSE50081 | 181 |
| Larsen_2007a | NA | GSE5828 | 59 |
| Yu_2008 | 18636107 | GSE5364 | 30 |
| Larsen_2007b | 17504995 | GSE5843 | 48 |
| Schabath_2016 | 26477306 | GSE72094 | 442 |
| Moriya_N |  | GSE7339 | 100 |
| BrolCt_2009 | 19176396;20810387 | GSE10445 | 72 |
| Noro_2017 | 27613525 | GSE74777 | 107 |
| Su_2007 | 17540040 | GSE7670 | 54 |
| Angulo_2008 | 17992665 | GSE8569 | 75 |
| Lee_2008 | 19010856 | GSE8894 | 138 |
| Mitra_2011 | 21242119 | GSE9971 | 27 |
| Beer_2002 | 12118244 |  | 96 |
| Bhattacharjee_2001 | 11707567 |  | 203 |
| Baty_2010 | 19833826 | GSE11117 | 44 |
| TCGA_LUAD_2016 | 25079552 |  | 576 |
| TCGA_LUSC_2016 | 22960745 |  | 552 |
| Takeuchi_2006 | 16549822;21465578 | GSE11969 | 163 |
| Xi_2008 | 18927117 | GSE12236 | 40 |
| Boelens_2011 | 20832896 | GSE12472 | 63 |

The availability and distribution of clinical variables across all studies are summarized in Figure 1 and Supplementary Table 1. The clinical variables we collected include tumor histology as defined by the 2015 WHO lung tumor classification system (Figure 2), as well as patient demographics, diagnosis, adjuvant therapy, smoking status, recurrence-free and overall survival time and status, and mutation status of some key cancer genes (Figure 1 and Supplementary Table 1).

### Quality control of data

During the construction of the lung cancer database, extensive efforts were made in the manual inspection of clinical data and quality control of expression data.

Specifically, manual data curation was done by examining the associated research paper and its supplementary files to check for consistency with the clinical data downloaded from GEO. We also looked for additional clinical information in the publication. In this study, substantial effort has been made to ensure the data quality, which is extremely important in utilizing public datasets. The following are a few examples of our manual curation from numerous instances: we checked if there were exclusion criteria in the paper that imposed restrictions on adjuvant therapy, tumor stage, etc.; when calculating the survival time we looked for surgical date, and if it was available we used it as the start date for survival time instead of the initial diagnosis date, since the gene expression data reflects the tumor profile on the surgical date; when certain samples were considered low quality and removed from analyses in the associated publication, we followed the same discretion to exclude such samples from our collection; we removed cell line samples to ensure our collection included exclusively patient samples; when tumor percentage information was available, we removed samples with less than 50% tumor content.

To perform quality control of the expression data input for meta-analysis, we implemented a method that checks for reproducibility across studies based on the concept of the integrative correlation coefficient (ICC)[24, 25]. The premise of this approach is that most of the pairwise gene-gene correlation should be preserved across different studies. The relationship of reproducibility between studies could be visualized by ICC-based clustering as shown in Supplementary Figure 3. When we calculated ICC, we considered that some gene-gene correlation could be tissue-type specific. Therefore, we further separated samples of different tissue types from the same study into distinct groups before we calculated the ICC. Indeed, in clusters defined by ICC, we found that in many cases, subgroups of different samples types from the same study do not cluster together; instead, samples of the same tissue type from different studies tend to cluster together. We identified a clade of 4 studies with very little correlation with other sample groups, and they were removed from subsequent meta-analyses. However, these studies are still available to use in the individual dataset-based analysis. On the other hand, the two RNA-seq datasets from TCGA revealed high correlation with datasets from microarray platforms, supporting the compatibility of datasets from different platforms based on our processing approach.

Finally, with access to qPCR measurement of 46 nuclear hormone receptor genes in 30 pairs of matched tumor and normal lung cancer samples, we were able to compare the standardized mean difference between tumor and normal tissue gene expression estimated from meta-analysis to the qPCR measurement results. A strong agreement was observed between the two results, supporting the validity of our meta-analysis and high quality of our datasets (Figure 3, Supplementary Table 4).

**Figure 3.**
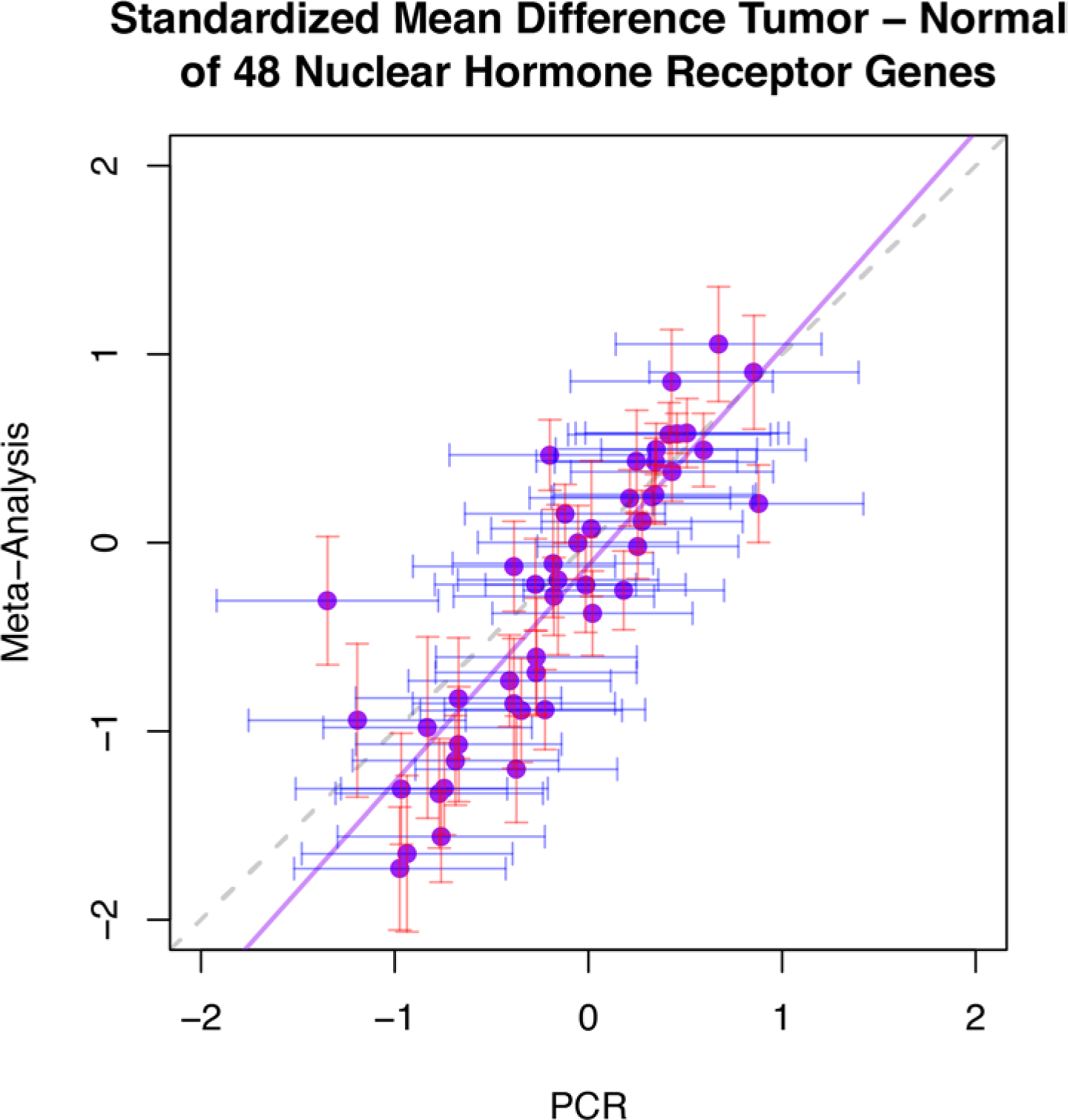
Meta-analysis estimates agree with qPCR measurements on tumor vs normal expression differences for 46 nuclear hormone receptor genes. Results from qPCR measurements of 30 tumor-normal pairs (x-axis values) and meta-analysis estimates from 21 studies (y-axis values) on gene expression differences between tumor and normal tissues for 46 nuclear hormone receptor genes were used to evaluate consistency between the two approaches. The values on the x-axis and y-axis are the standardized mean difference estimated by Hedges’ G method. The solid purple line represents a linear regression line, whereas the dashed gray line identifies where x equals y.

### The Lung Cancer Explorer (LCE)

Having established a high-quality lung cancer database, we constructed the user-friendly website Lung Cancer Explorer (LCE) (http://lce.biohpc.swmed.edu), allowing the cancer research community to gain easy access to our resources. Our dataset inventory and sources are described on the “DATA” page of LCE. The “ANALYSIS” page of LCE provides survival analysis, comparative analysis and co-expression analysis tools based on individual datasets, as well as meta-analysis tools based on multiple datasets. The functionality of these tools is described in detail in the following sections.

### Survival analysis in LCE

In LCE, survival analysis based on individual datasets is provided to allow users to assess the association between gene expression and prognosis. Users can separate the patient cohort into two groups by the expression level of the user-specified gene, and the survival differences between the two groups are reported in terms of hazard ratios and p values from log-rank test. Sometime, selecting a cutoff value to dichotomize continuous gene expression into two groups can be a tricky and ad hoc procedure. The LCE survival analysis module offers four options for cutoff value selection. Median gene expression level is used as the default cut-off option to define patient groups, as it usually provides the best statistical power by separating patients into two equal-sized groups. Other cutoff options implemented include “mean”, “cluster” and “custom”. In the Results panel, a Kaplan-Meier plot, table of summary statistics and Kernel density plot of the expression data is provided to the user. The density plot visualizes the distribution of the gene expression and facilitates the user to determine whether they should modify their choice of cut-off. For example, some genes might follow a bimodal distribution with two sub-populations of imbalanced sample sizes; under such a scenario the “cluster” option in cut-off selection would be a more rational choice than the default “median” option, as it separates the sample groups by a cut-off estimated from Gaussian mixture modelling assuming bimodal distribution. In Figure 4 we show some examples using genes *SMARCA4* and *KYNU* in two lung adenocarcinoma (ADC) studies. Bi-modal distribution of gene expression was observed in both cases (Figure 4a and d). *SMARCA4*, a well-known tumor suppressor gene [26], was under-expressed in a small fraction of samples from the Shedden_2008 study [27] and the corresponding patients have worse survival outcome (Figure 4c). In contrast, *KYNU* was over-expressed in a small proportion of samples in dataset Schabath_2016 [28] and the corresponding patients also have worse survival outcome. In both cases, results from survival analysis were more significant when the cut-off was selected by “cluster” as opposed to “median” (Figure 4b, c, e and f). With the built-in “cluster” option for cut-off selection, users can easily generate figures like Figure 4c and f and compare with the default “median” option like Figure 4b and e. In addition, we also allow the user to set a “custom” cut-off with their preferred value.

**Figure 4.**
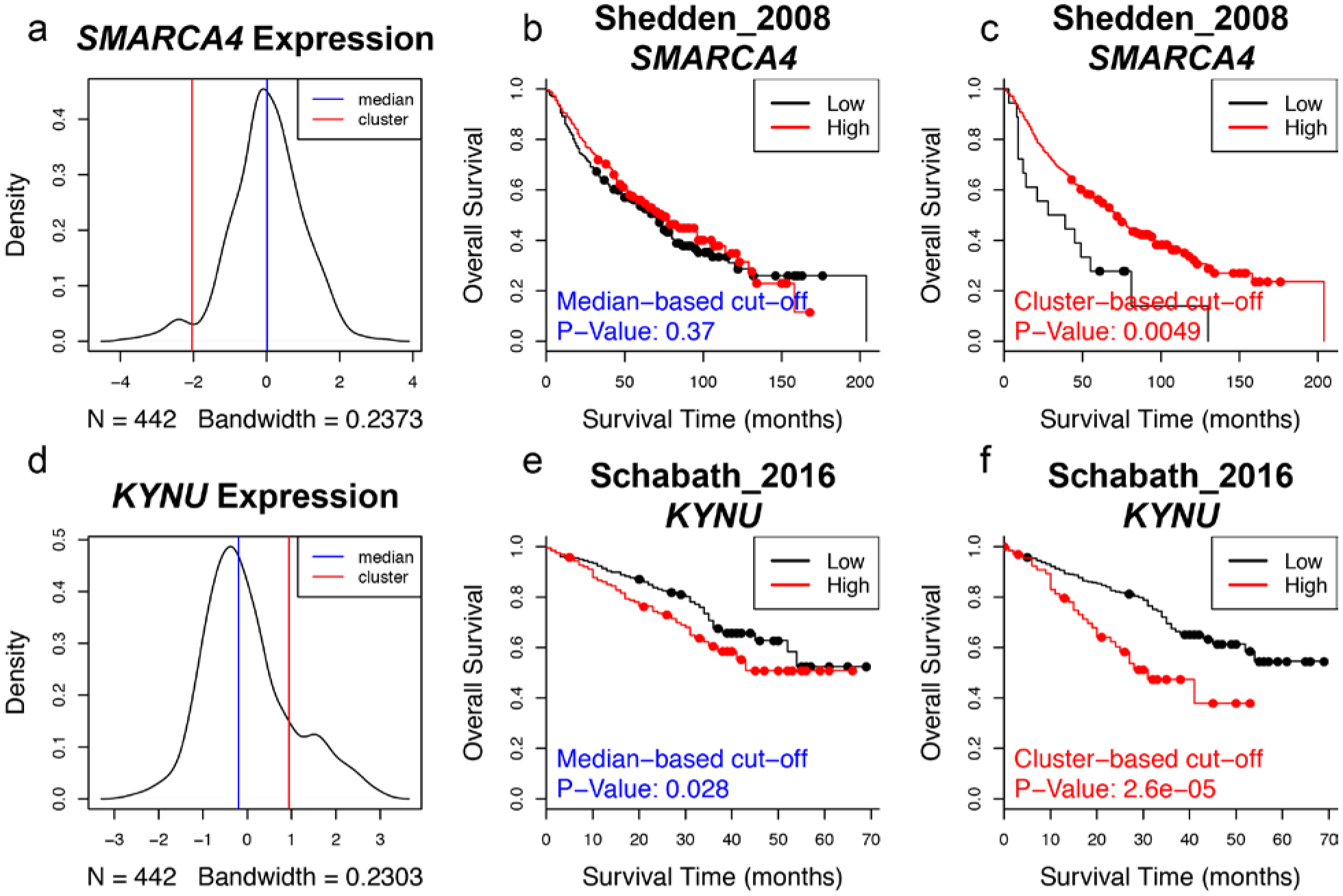
Examples of survival analysis with more significant results when cluster based cut-off is used. **a**, Bi-modal distribution of expression in Shedden_2008 dataset. The solid blue line marks the cut-off at the median, whereas the solid red line marks the cut-off determined by Gaussian mixture model. **b**, Kaplan Meier curves from the survival analysis of Shedden_2008 using groups defined by *SMARCA4* gene expression with cut-off at median. P-value from the log-rank test is denoted at the bottom left corner of the plot. **c**, Survival analysis of Shedden_2008 using groups defined by Gaussian mixture model of *SMARCA4* expression. **d**, Bi-modal distribution of *KYNU* expression in Schabath_2016 dataset. **e**, Survival analysis of Schabath_2016 using groups defined by *SMARCA4* gene expression with cut-off at median. **f**, Survival analysis of Schabath_2016 using groups defined by Gaussian mixture model of *KYNU* expression.

In the survival analysis, options are provided for users to select a group of patients by age, race, gender, smoking status and histology. This allows users to assess the association between the expression of a user-selected gene and patient survival (gene-survival association) within a user-defined subpopulation of patients. Supplementary Figure 4 provides some examples where we compare gene-survival association in different patient groups defined by genders. We show that for several studies, a stronger positive association between high klotho gene expression and overall survival could be observed in male patients as compared to female patients. Klotho, encoded by gene *KL*, is a well characterized anti-aging gene [29]. It has been observed that the extension of lifespan by klotho overexpression is more pronounced in males than in females [30], and only male but not female klotho mutant mice responded to a phosphorus restriction diet to extend lifespan [31]. In recent years, klotho has also been characterized as a tumor suppressor gene [32]. From our analyses, it is interesting to see that the tumor suppressing effect of klotho also seems to be higher in males than in females (Supplementary figure 4).

### Comparative analysis in LCE

Comparative analysis was implemented for users to assess the associations between a user-selected gene and clinical factors such as gender, age, histology types, disease stages, etc., within a specific dataset. The expression levels of the selected gene in the user-defined patient groups are shown in boxplots and p values of the expression differences are reported. In addition to group assignment based on a single clinical variable, a unique functionality of LCE is that users can define patient groups based on a combination of clinical factors. This provides a great extent of flexibility in hypothesis testing to understand the interactions between different clinical variables. For example, expression comparison of hemoglobin subunit delta encoding gene *HBD* in the TCGA_LUAD_2016 cohort shows that tumor samples have decreased *HBD* expression compared to normal samples (Supplementary Figure 5a), whereas samples from smokers and non-smokers have similar expression levels (Supplementary Figure 5d). However, by defining patient groups by both tissue type (tumor vs normal) and smoking status, we find the difference in *HBD* levels between normal and tumor tissues is significant only in smokers but not in non-smokers (Supplementary Figure 5b and c), and normal samples from smokers have elevated *HBD* expression compared to normal samples from non-smokers (Supplementary Figure 5f). In contrast, no difference in *HBD* expression was observed for tumor tissues from smokers vs non-smokers (Supplementary Figure 5b), nor do *HBD* expression levels differ in the tumor and normal tissues of non-smokers (Supplementary Figure 5f).

The results from these comparisons suggest that *HBD* expression is upregulated in normal lung tissue by smoking but is down-regulated again when tumors formed in smokers. We also observed similar trends in other hemoglobin subunit encoding genes *HBG1*, *HBG2* and *HBM,* which is consistent with the previous finding that hemoglobin levels increase in smokers [33].

### Meta-analysis in LCE

In LCE, meta-analysis tools are provided to allow users to address two questions: (1) differential expression between tumor and normal samples; and (2) survival association of gene expression. Meta-analysis not only provides users an overview of whether a significant trend could be consistently observed across multiple studies, it is also more powerful and precise than using any single dataset.

Results from both types of meta-analyses are visualized as forest plots. We provide three options, “All Cancer”, “Adenocarcinoma” (ADC), or “Squamous Cell Carcinoma” (SCC), to allow users to choose the type of studies they want to include for the meta-analysis since the survival association and expression difference between tumor and normal could be cancer-type specific.

For example, with lung cancer subtype specific meta-analysis, we found consistent down-regulation of RAR Related Orphan Receptor C (*RORC*) in multiple lung SCC studies (Figure 5B) but not in lung ADC studies (Figure 5A). Interestingly, *RORC* was also previously found in a 3-gene signature to distinguish lung ADC and lung SCC [34]. We also found that in multiple lung ADC studies, expression of Cell Division Cycle-Associated protein 2 (*CDCA2*) is associated with worse overall survival outcome (Figure 5C), whereas this trend was not observed for lung SCC datasets (Figure 5D).

**Figure 5.**
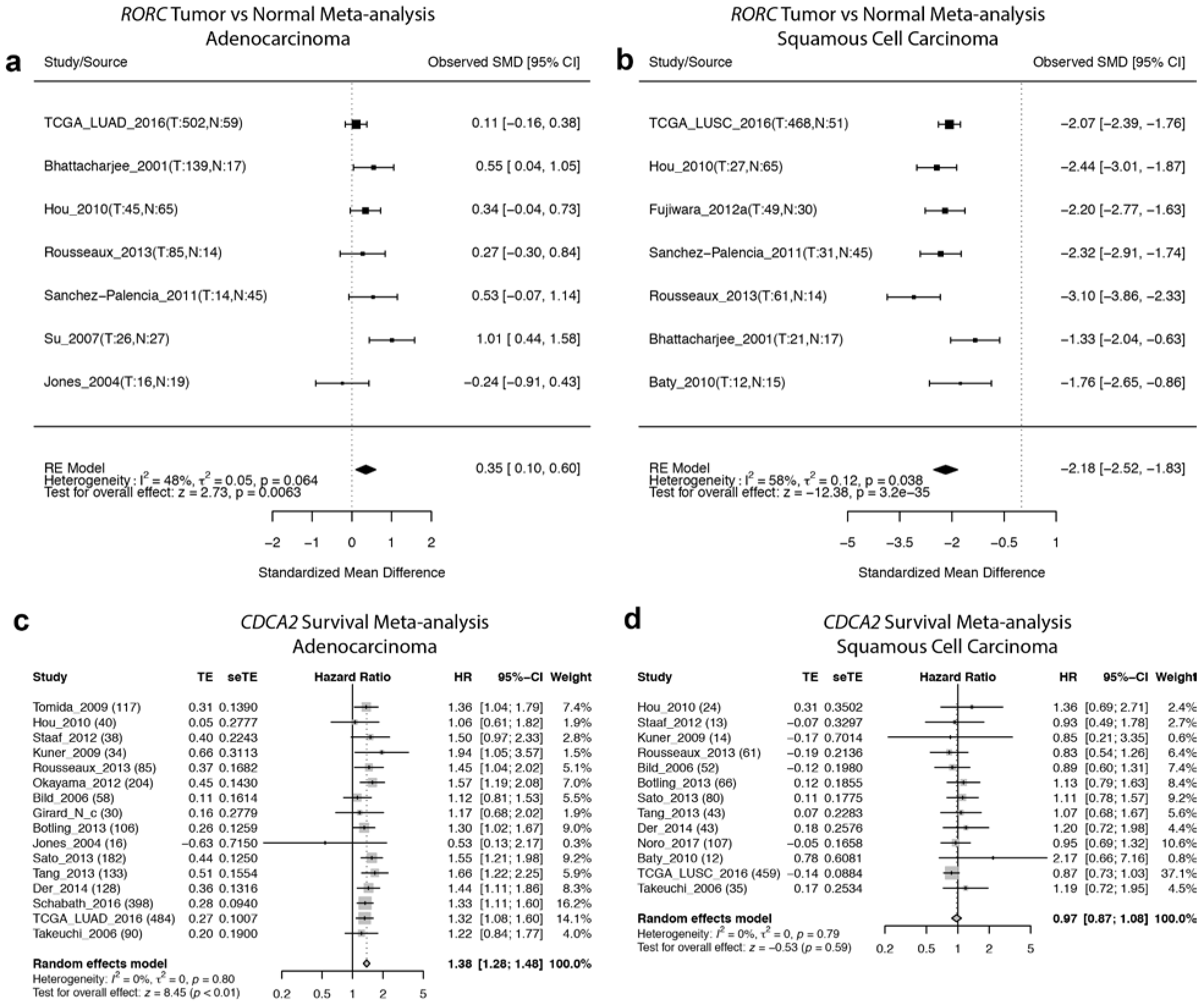
Examples of different meta-analysis results in lung adenocarcinoma vs squamous cell carcinoma. **a,b** *RORC* tumor vs normal meta-analyses in lung ADC studies (**a**) and lung SCC studies (**b**). **c,d** *CDCA2* survival meta-analyses in lung ADC studies (**a**) and lung SCC studies. Note that differential gene expression meta-analysis for *RORC* is only significant in lung SCC patients, whereas survival meta-analysis for *CDCA2* is only significant in lung ADC patients. In each forest plot, the name of each study is followed by the number of tumor and normal samples (tumor vs normal meta-analysis) or total tumor samples (survival meta-analysis). Abbreviations: SMD, standardized mean difference; TE, estimated treatment effect; seTE, standard error of treatment effect; HR, hazard ratio; CI, confidence interval.

Importantly, meta-analysis allows users to recognize the extent of reproducibility of a specific analysis across different datasets. In the forest plots generated by LCE meta-analysis module, we provide users with a heterogeneity test using the I^2^ statistic, which describes the percentage of variation across studies that is due to heterogeneity[35]. It is important to note that inconsistency of the results between different studies could arise from differences in patient population or sample procurement, as well as in data acquisition. In some cases, the results are more consistent for specific genes than others (Supplementary Figure 6). Hence, the meta-analysis tool provided by LCE allows users to identify discrepancies among different datasets in order to estimate the generalizability of the results.

### Correlation analysis in LCE

The correlation analysis tool from LCE provides users a heatmap to visualize the expression correlations among a list of user-defined genes in user-selected data sets. A high degree of expression correlation of genes often implies functional association, as genes involved in the same pathway or biological function are often subject to concerted regulation at transcription level [36]. Functional partners of the same gene could differ in a tissue-specific manner [37], and the gene network could also re-wire under a different disease context. In LCE we provide three options, “All”, “Lung Tumor” and “Normal”, to allow users to calculate a gene expression correlation matrix based on a specific sample type and subsequently generate a clustered heatmap, which conveniently allows users to identify changes in co-expression patterns of the user-defined gene list. One such example is provided in Figure 6, where we show that in tumor, there is a high degree of co-expression between Poly (ADP-ribose) polymerase-2 (*PARP2*) and 10 cell cycle genes (Figure 6a) selected from MSigDB “REACTOME_CELL_CYCLE” gene set [38, 39], whereas this co-expression is diminished in normal tissues (Figure 6b). This is consistent with the role of *PARP2* in DNA repair [40]; since genomic instability and mutation is a hallmark of cancer, the cancer specific co-expression of *PARP2* and cell cycle genes may indicate that *PARP2* is actively engaged in DNA repair while cancer cells divide. On the other hand, we found *PARP2* highly correlated with Zinc fingers C2H2-type genes (ZNF) [41] in normal but not cancer tissue (Figure 6 c and d). This normal specific co-expression of *PARP2* and ZNF genes may suggest alternative roles of *PARP2* in transcriptional regulation independent of its DNA repair function.

**Figure 6.**
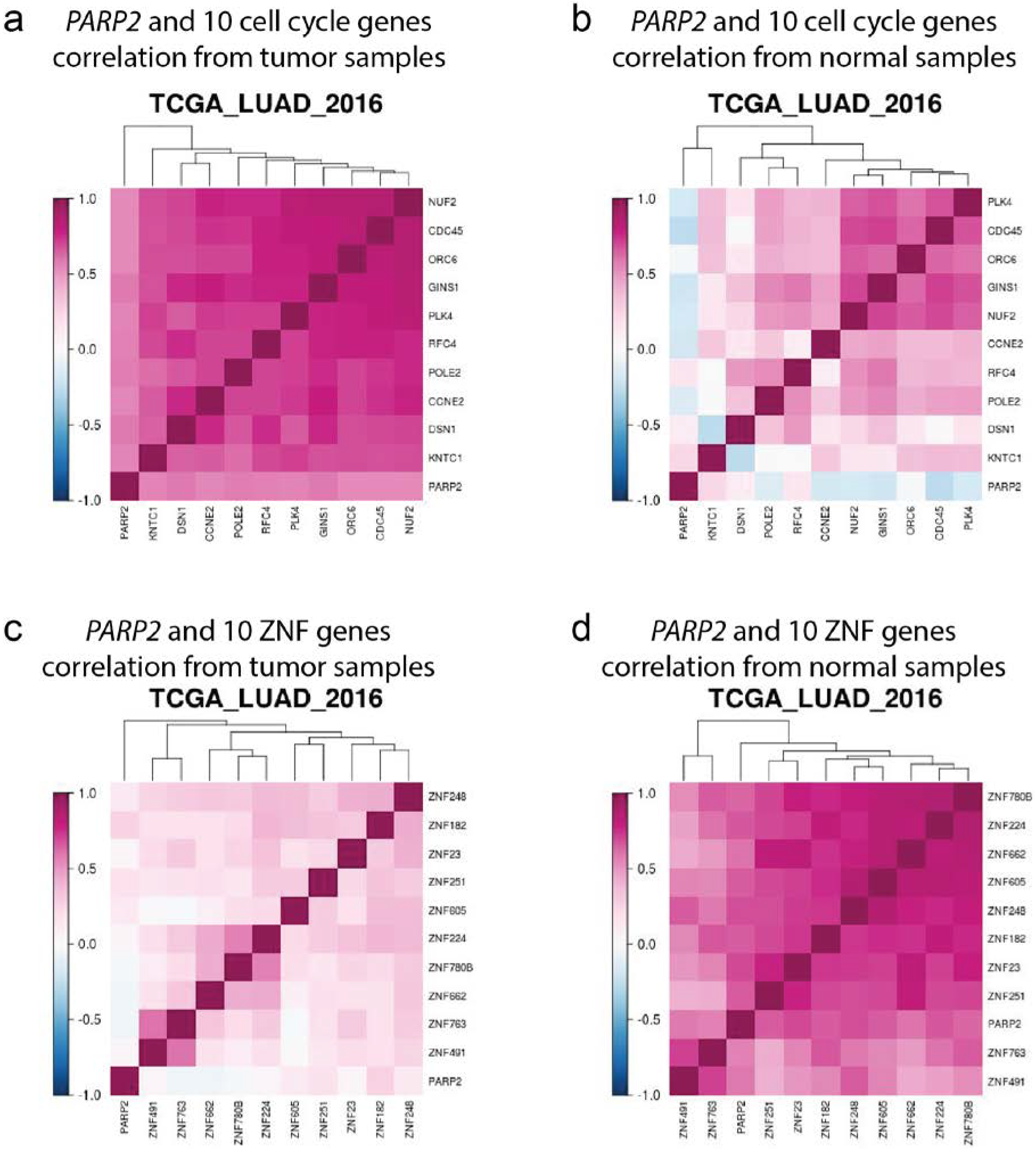
Different co-expression pattern between ***PARP2*** and cycle genes. **a,b,c,d** Heatmaps of gene-gene correlation matrices from TCGA_LUAD_2016 for *PARP2* and 10 selected cell cycle genes from tumor sample expression data (**a**) or normal sample expression data (**b**), and for *PARP2* and 10 selected C2H2 type zinc finger genes (ZNF) from tumor sample expression data (**c**) or normal sample expression data (**d**). The highly positive correlation between *PARP2* and cell cycle genes was seen only in tumor samples but not normal samples (**a** and **b**), whereas the high degree of positive correlation between *PARP2* and ZNF genes was observed only in normal tissue samples but not tumor samples (**c** and **d**).

## Discussion

In this paper, we described the construction of the LCE database for lung cancer gene expression analysis. It was carefully designed for lung cancer researchers to interrogate gene expression association with patient clinical features. As the collected datasets are highly heterogeneous, extensive efforts were put forth to reprocess and normalize expression data from 23 different expression profiling platforms, and a large amount of manual curation work was performed to standardize clinical terminology. Such manual inspection, though time consuming, greatly improves the data accuracy and usability, which sets our work apart from other databases. The resulting database with high-quality datasets enables versatile analysis tools in our Lung Cancer Explorer. We provide meta-analysis tools that summarize results across multiple datasets in the form of forest plots to allow users to gain a summary view of the overall trend and heterogeneity among studies. We also provide individual dataset-based analysis tools to allow users the flexibility to intricately formulate their analysis to best fit the research question. Results and biological insights we obtained from examples (Figures 4-6 and Supplementary Figures 4 and 5) demonstrated the unique advantages of our tools over the current publically available web tools, as none of these results could have been produced with the existing public tools.

We welcome users to contribute or suggest additional datasets to be evaluated and added to our lung cancer database. Suggestions can be made by leaving a comment at the contact page of LCE. It is in our plan to add a functionality to LCE to enable users to upload their own data to our database and perform analysis with our web application. In the future, we would also like to expand the lung cancer database to include cell line data and patient-derived xenograft (PDX) data. Besides gene expression data, other types of molecular profiling data (such as proteomic data, mutation data, copy number variation data, epigenomics data, microRNA data, etc.) and imaging data (such as H&E pathological slide images) will also be added to the lung cancer database. Separate data tables and supporting data dictionaries will be created for the new molecular data types. We will first identify studies within our collection that possess such data and add them to our database, then we will look for additional datasets that contain such molecular data as well as clinical data to add to our database. We will also expand the analysis tool repertoire on LCE to include multivariate analysis and other integrative analytical approaches.

Finally, we will conduct a variety of systematic analyses with the lung cancer database to generate testable hypothesis (for example, identification of genes associated with different oncogenotypes, gender, smoking status, etc. followed by gene set enrichment analysis). Results from such systematic analyses will be provided to the lung cancer research community to provoke hypothesis generation, testing and validation.

---

**Supplementary Figure 1.**
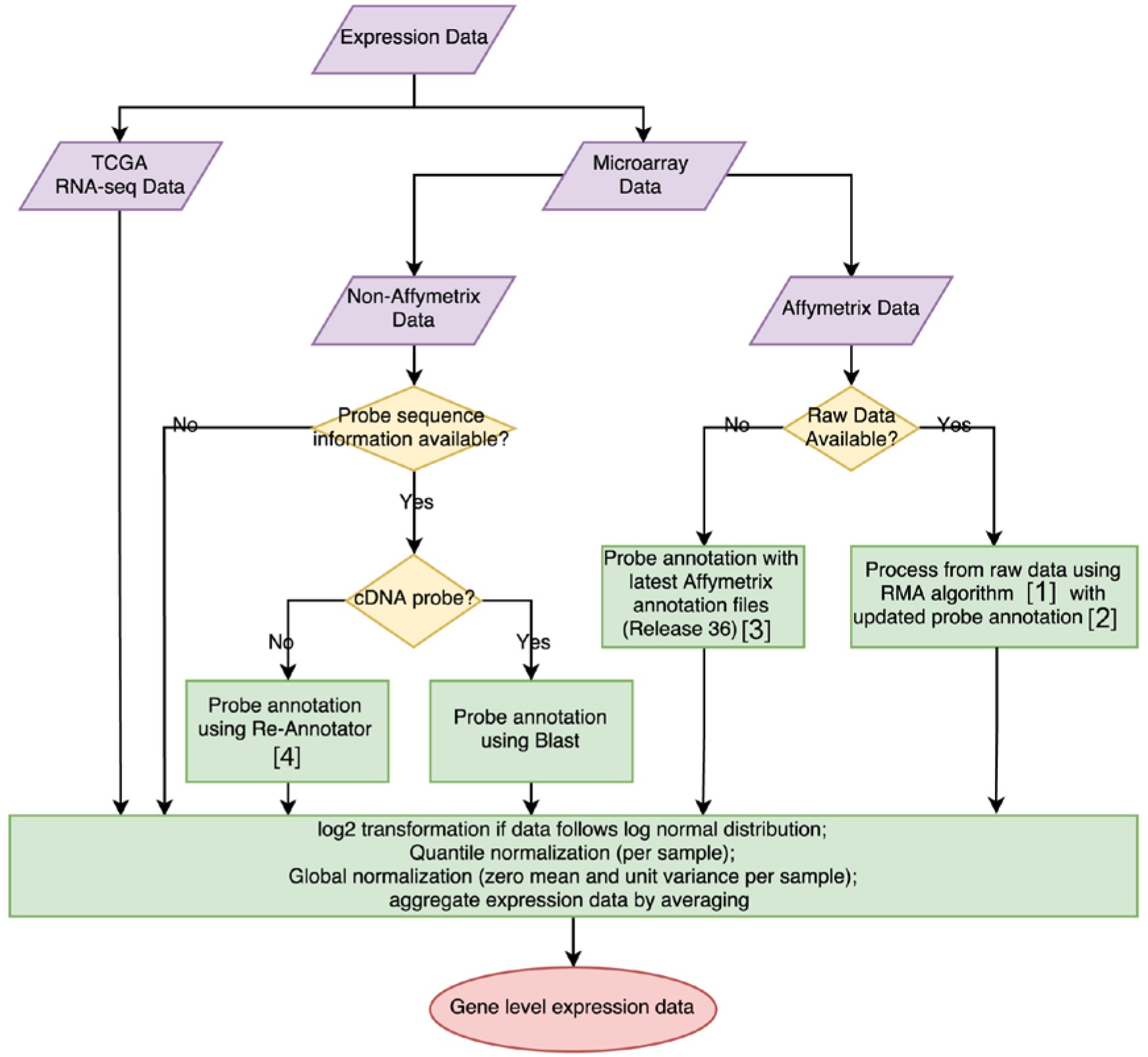
Flowchart of expression data processing strategies. Different strategies were adopted to process expression data from different platforms with or without raw data available for download. We re-annotated the probes whenever possible because the original probe design and annotations were often based on obsolete transcriptome databases. If the data were generated on an Affymetrix platform with raw data available for download, we processed from the raw data using an RMA algorithm [1] with updated probe annotation [2]; for data generated from the Affymetrix platform without raw data available for download, we mapped the probes to genes based on the most up-to-date probe annotation files (Release 36) downloaded from the Affymetrix website [3]. For data generated on a non-Affymetrix platform with probe sequence information available, re-annotation of datasets with short probes was performed using Re-Annotator [4], whereas datasets with long probes such as cDNA probes had probes remapped by Blast, similar to the strategy we adopted in ProbeMapper [5]. For data without probe sequence information available, we used the vender-provided probe annotation. Once the expression data have been mapped to gene level, we perform normalization steps for all datasets as follows: data was transformed if it followed a log normal distribution, quantile normalization and global normalization were performed subsequently, and the expression data was aggregated so that each sample has a unique expression value per gene.

**Supplementary Figure 2.**
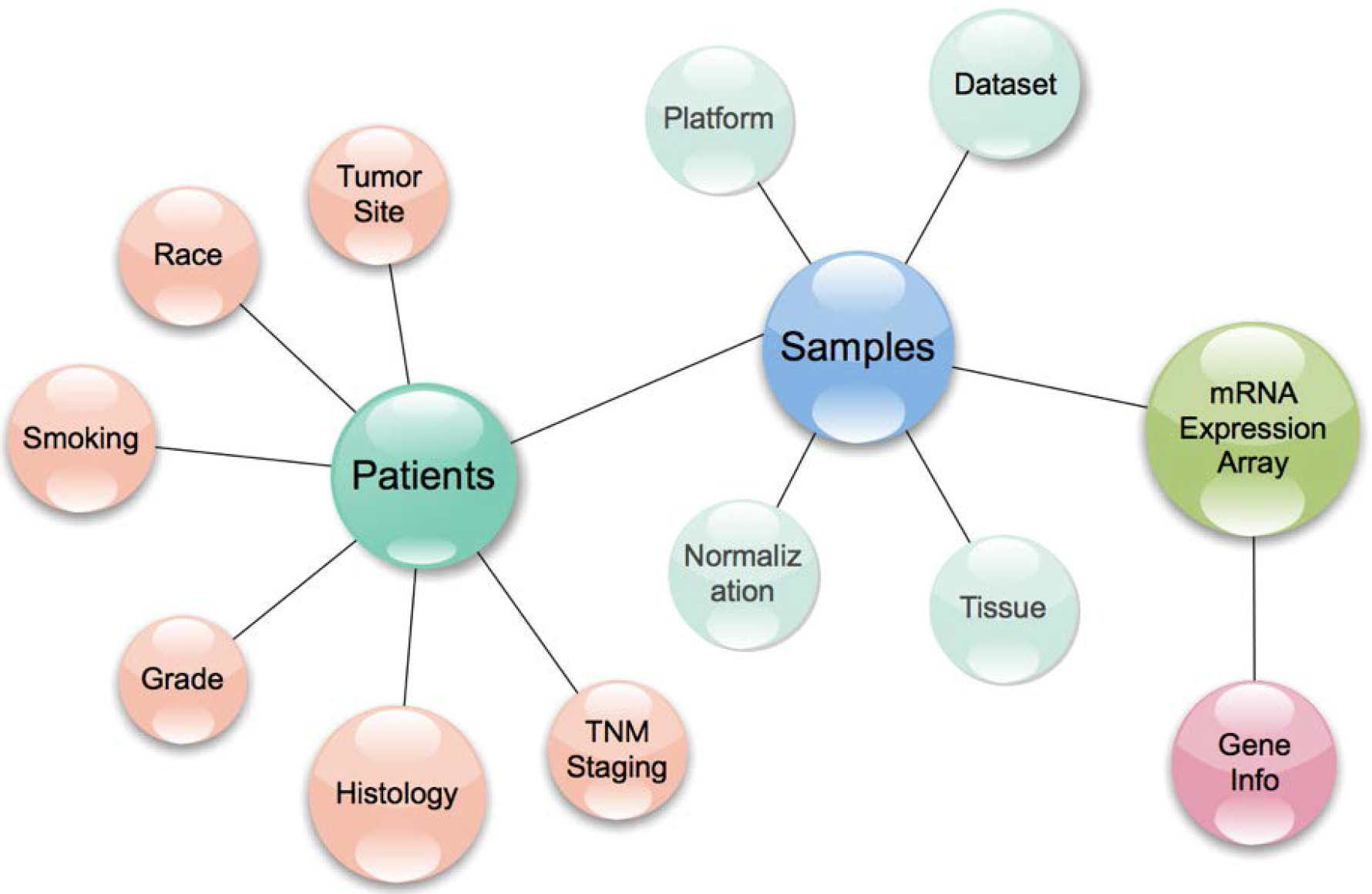
Structure of the lung cancer database. This schematic diagram shows the design of main data storage tables surrounded by supporting data dictionaries comprising our relational lung cancer database. Three main data storage tables are the patients table, samples table and the mRNA expression table. The patients table and the samples table are connected by patient ID, whereas the samples table and the mRNA expression table are connected by sample ID.

**Supplementary Figure 3.**
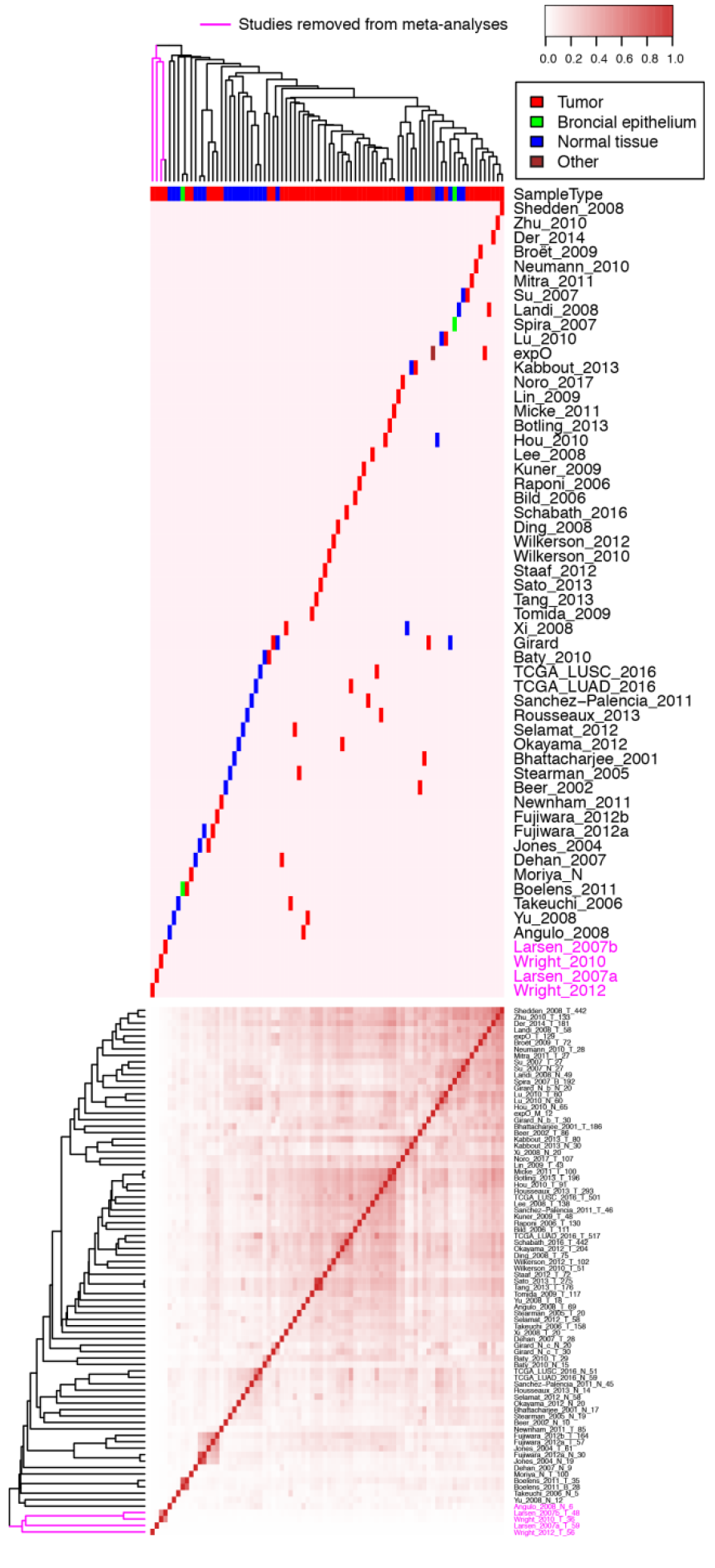
ICC Clustering of samples by sample type and sample source. 82 tissue-and-study-specific expression datasets were generated from 56 studies and were used to perform quality control of expression datasets used for meta-analysis. Integrative correlation coefficients were calculated based on the global expression correlation among these 82 expression data sets and were visualized in the heatmap. Datasets were clustered by hierarchical clustering using the average linkage method. Column-side color labels of the heatmap highlight the location of datasets originating from specific studies, and different tissue types were represented by different colors. Four studies that had little ICC with all other studies were excluded from meta-analysis in LCE.

**Supplementary Figure 4.**
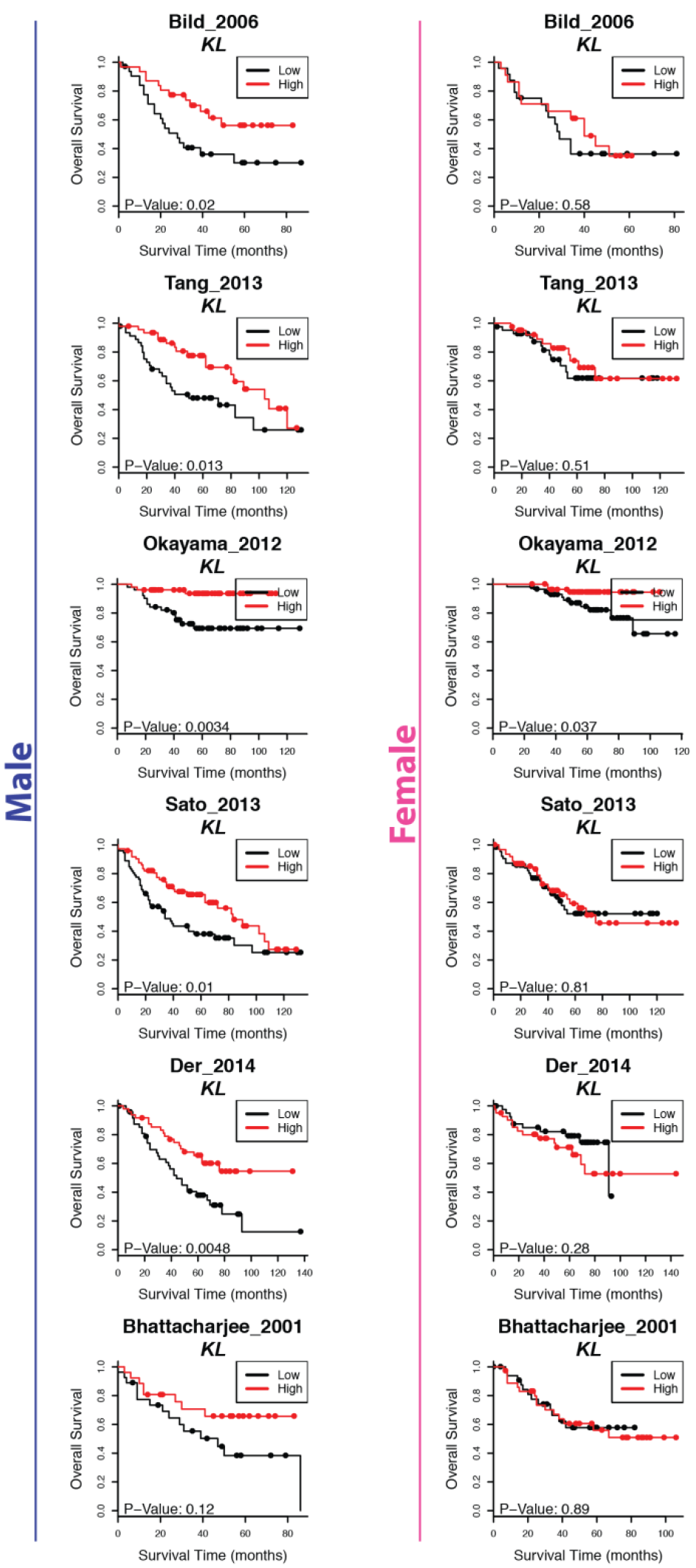
High klotho expression has more significant association with positive survival outcome in males. For each of the six selected studies, survival analysis assessing prognosis association of KL gene expression was performed for male patients or female patients only. In each analysis, the median was used as a cut-off for dichotomizing patients. In all six studies, a more significant association with better prognosis was found in the male patients compared to the female patients.

**Supplementary Figure 5.**
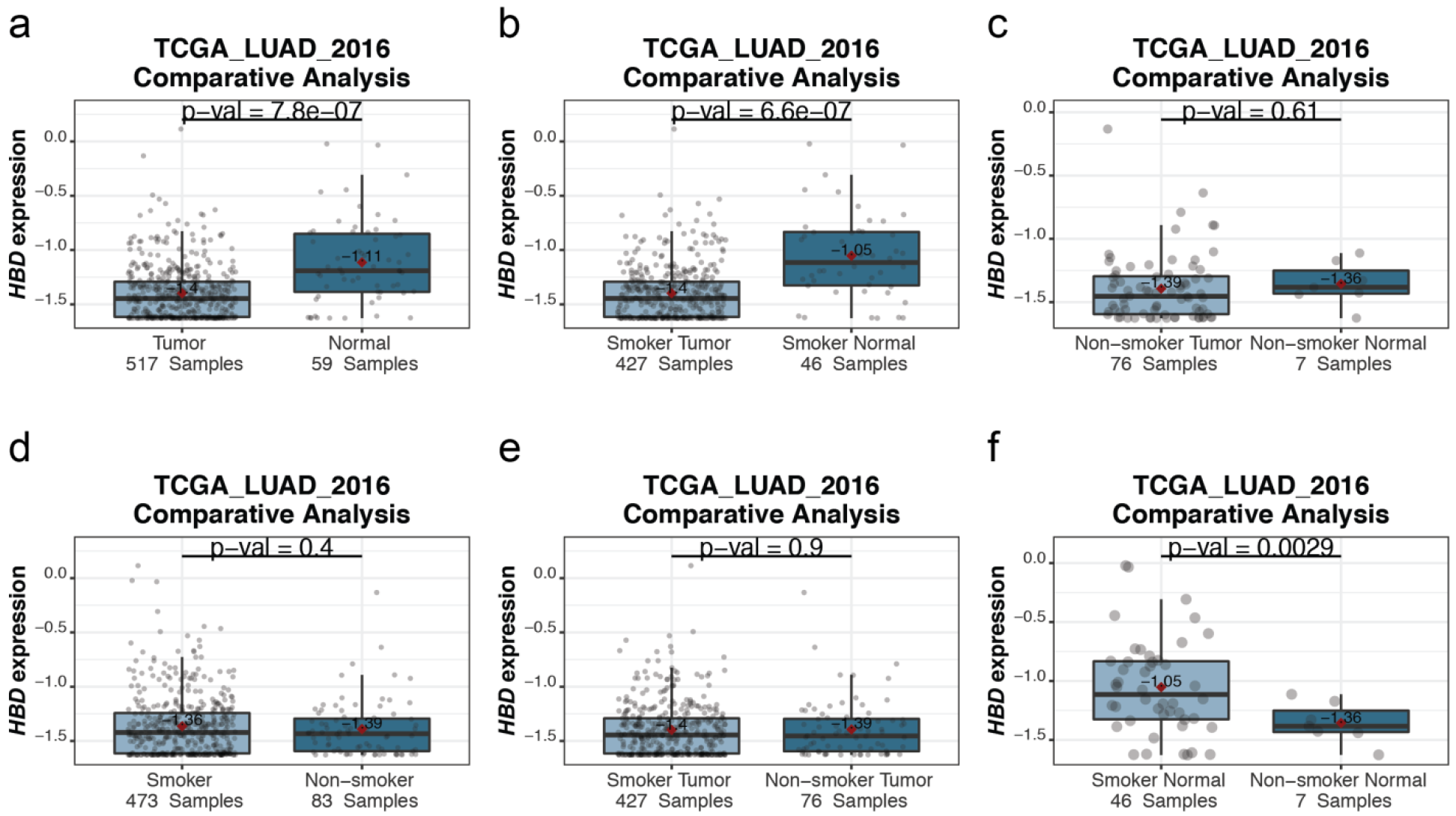
Interaction between sample tissue type and smoking status in HBD gene expression. **a,d** Boxplots comparing *HBD* gene expression between two groups dichotomized on a single clinical variable: tissue type (**a**) or smoking status (**d**). **b,c,e,f** Boxplots comparing *HBD* gene expression between two groups defined by a combination of two clinical variables: different tissues in smoker (**b**), different tissues in non-smoker (**c**), tumor from patients with different smoking status (**e**) and normal tissues from patients with different smoking status (**f**).

**Supplementary Figure 6.**
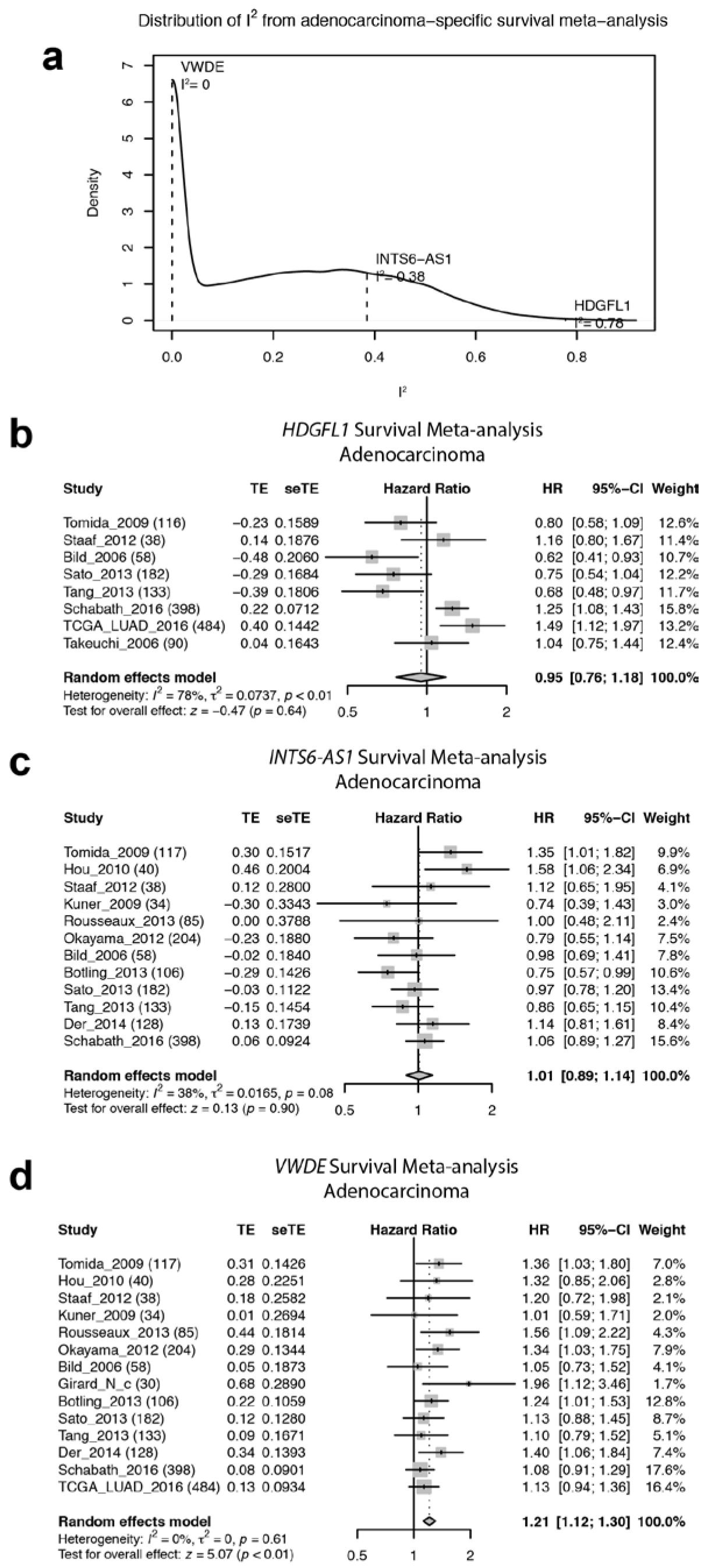
Adenocarcinoma meta-analysis of survival-gene expression association. **a** Density estimation of I^2^ distribution. Three genes with different I^2^ statistics were selected as examples in b, c and d. A larger I^2^ value suggests a larger degree of heterogeneity across studies, whereas a smaller I^2^ value is reflective of a higher degree of consistency among studies. **b,c,d** Example forest plots of survival meta-analysis with different heterogeneity: large (**b**), intermediate (**c**) and small (**d**).

